# Evaluation of strategies for the assembly of diverse bacterial genomes using MinION long-read sequencing

**DOI:** 10.1101/362673

**Authors:** Sarah Goldstein, Lidia Beka, Joerg Graf, Jonathan L. Klassen

## Abstract

**Background:** Short-read sequencing technologies have made microbial genome sequencing cheap and accessible. However, closing genomes is often costly and assembling short reads from genomes that are repetitive and/or have extreme %GC content remains challenging. Long-read, single-molecule sequencing technologies such as the Oxford Nanopore MinION have the potential to overcome these difficulties, although the best approach for harnessing their potential remains poorly evaluated.

**Results:** We sequenced nine bacterial genomes spanning a wide range of GC contents using Illumina MiSeq and Oxford Nanopore MinION sequencing technologies to determine the advantages of each approach, both individually and combined. Assemblies using only MiSeq reads were highly accurate but lacked contiguity, a deficiency that was partially overcome by adding MinION reads to these assemblies. Even more contiguous genome assemblies were generated by using MinION reads for initial assembly, but these were more error-prone and required further polishing. This was especially pronounced when Illumina libraries were biased, as was the case for our strains with both high and low GC content. Increased genome contiguity dramatically improved the annotation of insertion sequences and secondary metabolite biosynthetic gene clusters, likely because long-reads can disambiguate these highly repetitive but biologically important genomic regions.

**Conclusions:** Genome assembly using short-reads is challenged by repetitive sequences and extreme GC contents. Our results indicate that these difficulties can be largely overcome by using single-molecule, long-read sequencing technologies such as the Oxford Nanopore MinION. Using MinION reads for assembly followed by polishing with Illumina reads generated the most contiguous genomes and enabled the accurate annotation of important but difficult to sequence genomic features such as insertion sequences and secondary metabolite biosynthetic gene clusters. The combination of Oxford Nanopore and Illumina sequencing is cost effective and dramatically advances studies of microbial evolution and genome-driven drug discovery.

## Introduction

Microbial genome sequencing has revealed how microorganisms adapt, evolve, and contribute to health and disease [1, 2]. Although these were once enterprise-level projects, technological advances have now reached the point where microbial genomes can be sequenced routinely by small teams for a few hundred dollars [1]. These advances have particularly been driven by the maturation of short-read sequencing technologies such as those marketed by Illumina, which generate highly accurate reads (>99%) with lengths ranging from 75-300bp [1]. Although Illumina technologies currently dominate the sequencing market [1, 2], difficulties remain that require further technological advances to fully realize the potential of microbial genome sequencing.

By their very nature, short reads alone cannot disambiguate repetitive genomic regions that are longer than their read length. Unfortunately, such repetitive regions are common in microbial genomes [3–6], and include ribosomal genes, transposons, insertion sequences, CRISPR arrays, rhs toxins, secondary metabolite biosynthetic gene clusters, and many others [5]. Repeats lead to unresolvable loops in the underlying genome assembly graph that are ultimately fragmented into contigs [5, 7]. Because of this, short reads are theoretically incapable of closing most microbial genomes.

Genome assembly using most short-read datasets is also challenged by biases that occur during library preparation and that cause some genomic regions to be excluded from sequencing libraries. Common short-read library preparation methods (e.g., the Illumina Nextera protocol) include PCR amplification steps that are biased against regions of the genome with extreme GC contents [8–12]. Such regions are common among bacteria, whose average GC content ranges widely from 25% to 75% [13]. Library preparation protocols that use transposases to fragment DNA may also non-randomly shear genomes during library preparation [14], causing further biases that limit the utility of short-read sequencing.

De novo genome assembly algorithms struggle to assemble genomes when intergenic repeats are present and GC biases skew sequencing coverage [15, 16]. Fragmentation of such genomes prevents the accurate identification of mobile elements, the detection of horizontal gene transfers, the determination of gene copy number, and the discovery of biotechnologically important gene clusters such as those that encode for the production of secondary metabolites [16, 17]. These deficiencies significantly lower the informational value of draft-quality genomes [18, 19].

Recently, long-read, single-molecule sequencing has overcome some of the deficiencies of short-read sequencing. Library preparation protocols for single-molecule sequencing typically avoid bias-prone PCR steps, and long read lengths span genomic repeats to unambiguously resolve complex genomic regions. Some Illumina-based technologies such as mate pair libraries and linked reads (e.g., as commercialized by 10X Genomics) can also generate positionally linked sequences that span complex genomic repeats [1], but these technologies still require library preparation protocols that are subject to the biases discussed above. Pacific Biosciences (PacBio) currently markets the most widely used single-molecule sequencing technology, which can produce > 7 Gb per run with read lengths averaging >12 kbp [1]. Although the error rate for PacBio sequencing is high (~13%), these errors are near-randomly distributed and can largely be corrected during assembly with adequate sequencing coverage [7]. Unlike some Illumina sequencers (e.g., the MiSeq and MiniSeq), all PacBio sequencers require considerable capital investment, limiting general access to these technologies in individual laboratories. Nevertheless, PacBio sequencing has shown the enormous potential for long-read, single-molecule sequencing to routinely produce high-quality microbial genome assemblies that overcome many of the deficiencies of short-read sequencing.

The Oxford Nanopore Technologies (ONT) MinION is a more recently developed long-read, single-molecule sequencing instrument. The MinION is a small benchtop device that can plug directly into a laptop via a USB3 port [20] and that requires a relatively small upfront financial investment relative to PacBio instruments [1]. This affordability and simplicity has enabled the rapid uptake of MinION sequencing by individual labs worldwide, and facilitated new applications such as tracking disease outbreaks in low-resource environments [21]. MinION read lengths have no theoretical limit and reads >2 million bp long have been reported [22]. As with PacBio, MinION read quality is low compared to short read sequencing technologies [23, 24]. These errors are less randomly distributed than for PacBio sequencing [25], meaning that increased read depth alone cannot completely overcome this high error rate, at least currently. However, error rates and bias profiles are expected to improve as the MinION and its associated base-calling software continues to develop, e.g., as demonstrated by the increased accuracy of the new Scrappy base caller that is currently under development by ONT [26].

Two main strategies have been used to assemble bacterial genomes using MinION sequencing [27, 28]. In the first, MinION reads are used to enhance genome assemblies that are generated from short-read Illumina data. Here, MinION reads can scaffold contigs generated by Illumina sequencing [29–31] or be directly used in the assembly process to disambiguate regions of the assembly graph that cannot be resolved by Illumina sequencing alone (e.g., as implemented in the popular SPAdes and Unicycler software [32, 33]). Alternatively, MinION reads alone are used to generate an initial genome assembly [34, 35] that can then be further polished using either MinION or Illumina reads [34, 36]. Such polishing is highly recommended for MinION-based genome assemblies due to their higher error rates relative to assemblies based on Illumina data [17, 26, 27, 37, 38]. The increasing maturity and throughput of MinION sequencing is leading to its adoption for routine microbial genome sequencing [39–41].

Both MinION-only [34, 35] and Illumina-hybrid methods [32, 33] have been validated extensively for bacteria with low and average GC contents. However, whether these approaches offer advantages when assembling bacterial genomes with high GC content remains unclear [42] (but see [43]). We therefore compared the ability of Illumina and MinION sequencing technologies to produce high-quality assemblies of genomes from three bacterial genera (Flavobacterium, Aeromonas, and Pseudonocardia) that range in GC content from 31-73% (Table 1). Flavobacterium spp. are gliding bacteria that can be found in diverse environments and that include important fish pathogens. Aeromonas spp. are ubiquitous in aquatic environments and can cause diseases in humans and fish or form beneficial symbioses, e.g., with fish and leeches [44]. Pseudonocardia sp. are members of the Actinobacteria and, like many other members of this class, are important producers of antibiotics such as those involved in defensive symbioses with ants (e.g., [45]). Our results validate MinION sequencing’s ability to generate high-quality assemblies for all of these genomes, and especially emphasize the advantages of MinION sequencing when unbiased Illumina sequencing libraries are difficult to generate, e.g., for Actinobacteria with high GC content. These improved genome assemblies dramatically improve the annotation of repetitive genomic regions such as insertion sequences and secondary metabolite biosynthetic gene clusters (BGCs). MinION sequencing therefore has strong potential to overcome current limitations of short-read sequencing technologies and catalyze improved understanding of genome evolution and exploitation of genomic data for drug discovery.

**Table 1.** Bacteria used in this study

| Strain ID | Phylum | Genus | Species | % GC Content | Expected Genome Size (Mbps) |
| --- | --- | --- | --- | --- | --- |
| ***Ps* JKS002128** | Actinobacteria | *Pseudonocardia* | *sp* | 73.12 | 6.60 |
| ***Ps* JKS002072** | Actinobacteria | *Pseudonocardia* | *sp* | 73.69 | 6.21 |
| ***Ps* JKS002056** | Actinobacteria | *Pseudonocardia* | *sp* | 73.31 | 6.54 |
| ***Av* JG3** | Proteobacteria | *Aeromonas* | *veronii* | 58.64 | 4.49 |
| ***Av* CIP107763<sup>T</sup>** | Proteobacteria | *Aeromonas* | *culicicola*<sup>a</sup> | 58.80 | 4.34 |
| ***Ah* CA-13-1** | Proteobacteria | *Aeromonas* | *hydrophila* | 61.29 | 4.76 |
| ***Fs* ARS-166-14** | Bacteroidetes | *Flavobacterium* | *sp* | 31.61 | 3.31 |
| ***Fc* FC-100715-19** | Bacteroidetes | *Flavobacterium* | *columnare* | 31.59 | 3.32 |
| ***Fc* FC-08-1215-1** | Bacteroidetes | *Flavobacterium* | *columnare* | 31.56 | 3.31 |

<sup>a</sup>CIP107763<sup>T</sup> is the type strain for *Aeromonas culicicola*, which is a later subjective synonym of *A. veronii*.

## Methods

### Description of Strains

Three Aeromonas strains were used in this study. Aeromonas hydrophila str. CA-13-1 (hereafter Ah CA-13-1) was isolated from the wound of a patient undergoing post-operative leech therapy in 2013 [46]. Aeromonas veronii str. CIP107763^T^ (hereafter Av CIP107763^T^) was isolated from a mosquito midgut in France in 2015 and sequenced previously [47]. A. veronii str. JG3 (hereafter Av JG3) is a derivative of a medicinal leech isolate Hm21 [48]. All Aeromonas strains were grown either in LB broth or on agar plates for 16 hours at 30°C.

The Flavobacterium strains used in this study were all isolated from necrotic gill tissues of farmed rainbow trout, Onchorhyncus mykiss. Flavobacterium sp. str. ARS-166-14 (hereafter Fs ARS-166-14) was isolated in October 2014, Flavobacterium columnare str. FC-081215-1 (hereafter Fc FC-081215-1) was isolated in August 2015, and F. columnare str. FC-100715-19 (hereafter Fc FC-100715-19) was isolated in October 2015, all on TYES agar. Frozen cells were grown on TYES agar, incubated for three days at 20°C, and then grown in liquid TYES broth for another 3 days at 15°C for Fs ARS-166-14 and 25°C for Fc FC-100715-19 and Fc FC-08-1215-1 [49].

The *Pseudonocardia* bacteria sequenced during this study were isolated from individual *Trachymyrmex septentrionalis* ants collected from three locations within the United States: Paynes Creek Historic State Park, FL (*Pseudonocardia* sp. str. JKS002056, hereafter Ps JKS002056), Magnolia Springs State Park, GA (*Pseudonocardia* sp. str. JKS002072, hereafter Ps JKS002072), and Jones Lake State Park, NC (*Pseudonocardia* sp. str. Ps JKS002128). *Pseudonocardia* were visible as white patches on the ants’ propleural plates, which were scraped using a sterile needle under a dissecting microscope to isolate *Pseudonocardia* following Marsh [50].

### DNA Isolation

DNA was extracted from *Aeromonas and Flavobacterium* isolates following a modified version of a previously published protocol for large scale genomic DNA isolation [51, 52]. DNA in solution was not micropipetted during these extractions to minimize DNA fragmentation. DNA was extracted from single *Pseudonocardia* colonies using the Epicentre MasterPure Complete DNA and RNA kit following the manufacture’s protocol. Each *Pseudonocardia* extraction was performed in triplicate using wide bore tips and taking care to pipette slowly to prevent DNA shearing.

### Library Preparation and Sequencing

The quality of all extracted DNA was assessed using an Agilent TapeStation 2200 protocol for genomic DNA, an Agilent 2100 Bioanalyzer (High sensitivity DNA chip), and/or a Nanodrop spectrophotometer. All libraries were quantified using a Qubit^®^ 2.0 fluorometer. For the *Aeromonas and Flavobacterium* strains, NexteraXT Illumina sequences were constructed by following the manufacturer’s instructions for genomic tagmentation, PCR of tagged DNA, and PCR product cleanup. Libraries were diluted to 4nM for loading onto an Illumina MiSeq. TruSeq DNA PCR-Free libraries were created for each *Pseudonocardia* strain following the manufacturer’s protocol, shearing the DNA to 550 bp fragments using a Covaris M22 Focused-ultrasonicator. All Illumina libraries were sequenced on an Illumina MiSeq using the 2×250bp protocol at the University of Connecticut Microbial Analysis Research and Services (MARS) facility. Demultiplexing was performed using Illumina Basespace (https://basespace.illumina.com/home/index).

All genomes were also sequenced on a MK1B MinION device using R9.4 flow cells. *Aeromonas and Flavobacterium* libraries were prepared using the ONT EXP-NBD103 Barcode kit and the ONT “Native Barcoding Genomic DNA Sequencing for the MinION Device” protocol (downloaded from https://nanoporetech.com/resource-centre/protocols on Oct 20, 2017) and performed without optional shearing steps to select for long reads. *Pseudonocardia* libraries were prepared using the ONT “1D gDNA Selecting for Long Reads Using SQK-LSK108” protocol (downloaded from https://nanoporetech.com/resource-centre/protocols on Dec 20, 2016) All strains were sequenced using the ONT MinKNOW NC_48h_Sequencing_Run_FLO-MIN107_SQKLSK108 protocol, except for JKS002056, which was sequenced using the older NC_48h_Sequencing_Run_FLO-MIN106_SQK-LSK108 MinKNOW protocol. The run duration ranged from 12 to 48 hours. Strains Av JG3, Fc FC-100715-19, and Ps JKS002072 were sequenced using two separate MinION runs that were combined for all analyses, except for the Av JG3 Canu+Nanopolish assembly where the few MinION reads (<3000) from the first run were excluded because of their being processed using base calling software that was incompatible with Nanopolish.

### Base calling and Read Preparation

MinION reads for Ps JKS002056 and the first Av JG3 run were base-called using the ONT Metrichor 1D protocol and locally using MinKNOW (ONT; Oct 20, 2017 release) respectively. All other MinION reads were based-called using Albacore (v.1.2.4). These software choices were determined by changes made by ONT to their cloud-based base calling system. All raw data was deposited in the NCBI database under the BioProject number PRJNA477342.

We assessed Illumina read quality using FastQC (v.0.11.5, available from http://www.bioinformatics.babraham.ac.uk/projects/fastqc/). Trimmomatic (v.0.36; [53]) was used remove Illumina adapters, bases at each end of the read with an average Phred score <15 over a 4 bp window, and reads ≤36 basepairs long. Poretools version 0.6.0 [54] was used to assess the quality of each MinION dataset and to generate fastq files from basecalled fast5 files. Barcodes and reads that contained an internal barcode adapter sequence were removed using Porechop version 0.2.3 (available from https://github.com/rrwick/Porechop). Nanofilt (v.1.0.5, available from, https://github.com/wdecoster/nanofilt) was used to remove reads shorter than 500 basepairs or having an average quality score <9.

### Genome Assembly

We used several approaches to construct de novo assemblies of each genome. First, we constructed MiSeq-only short read assemblies using SPAdes (v.3.11.1) [33] and Unicycler (v.0.4.3) [32] (v.0.4.3), representing the current state of the art. Second, we added MinION reads to these MiSeq-based assemblies to disambiguate ambiguous regions in the MiSeq sequencing graph, creating SPAdes-hybrid and Unicycler-hybrid assemblies. Third, we constructed MinION-only long-read assemblies using Canu (v.1.5) [35]. These MinION-only Canu assemblies were polished using the same MinION reads to create Canu+Nanopolish assemblies by aligning MinION reads to the Canu assembly using BWA (v.0.7.15) [55] and Samtools (v.1.3.1) [56], and then using Nanopolish (v.3.2.5) [34] for assembly polishing. A second iteration of Nanopolish was completed for strain Ps JKS002128 but did not significantly improve its accuracy (data not shown), and so this strategy was not pursued further. The Canu assemblies were alternatively polished using MiSeq reads to create Canu+Pilon assemblies. MiSeq reads were aligned to the Canu genome using BWA (v.0.7.15) and Samtools (v.1.3.1) and then Pilon (v.1.22) [36] was used for assembly polishing. In total, we created seven assemblies for each genome: four based primarily on MiSeq data (SPAdes, Unicycler, SPAdes-hybrid, and Unicycler-hybrid) and three based primarily on MinION data (Canu, Canu+Nanopolish, Canu+Pilon). All commands used for the computational analyses in this study are included in the Supplementary Material.

### Depth of Coverage

MinION data was subsampled from Av JG3, Fs ARS-166-14, and Ps JKS002128 to determine the minimum read depth required to create contiguous MinION-based assemblies. Fast5-formatted reads for each strain were subsampled in the order that they were acquired from the MinION sequencer to achieve 10X, 20X, 30X, 40X, 50X, (for Fs ARS-166-14, Av JG3 and Ps JKS002128), 60X (Fs ARS-166-14 and Ps JKS002128 only), and 70X (Ps JKS002128 only) coverage of the Canu assembly for each strain, calculated using the mean MinION read length for each strain (Table 2). This strategy was used to simulate runs stopped after achieving each level of coverage. All data was processed and assembled using Canu as described above.

**Table 2.**
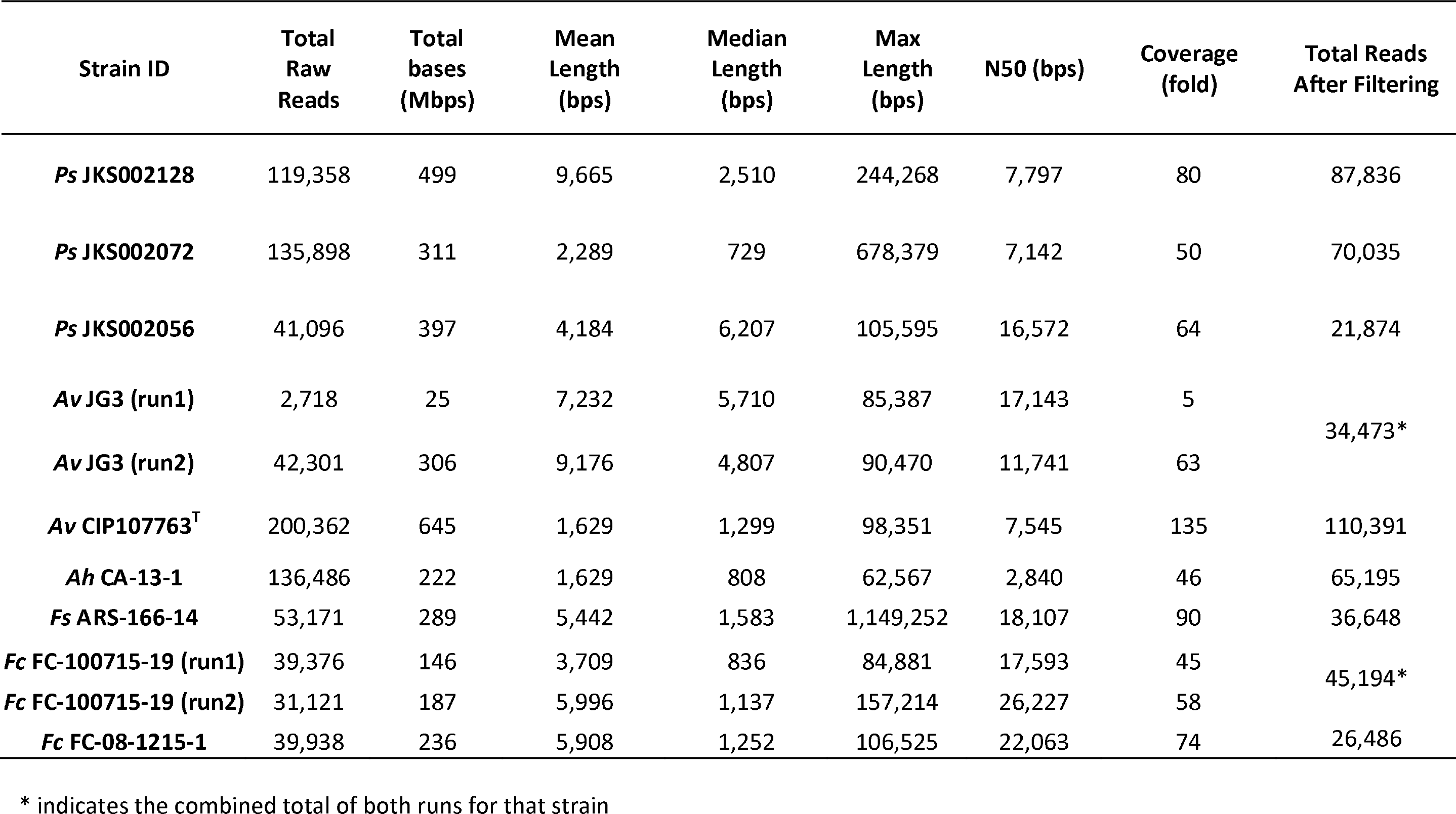
Summary of MinION sequencing

| Strain ID | Total Raw Reads | Total bases (Mbps) | Mean Length (bps) | Median Length (bps) | Max Length (bps) | N50 (bps) | Coverage (fold) | Total Reads After Filtering |
| --- | --- | --- | --- | --- | --- | --- | --- | --- |
| *Ps* JKS002128 | 119,358 | 499 | 9,665 | 2,510 | 244,268 | 7,797 | 80 | 87,836 |
| *Ps* JKS002072 | 135,898 | 311 | 2,289 | 729 | 678,379 | 7,142 | 50 | 70,035 |
| *Ps* JKS002056 | 41,096 | 397 | 4,184 | 6,207 | 105,595 | 16,572 | 64 | 21,874 |
| *Av* JG3 (run1) | 2,718 | 25 | 7,232 | 5,710 | 85,387 | 17,143 | 5 | 34,473* |
| *Av* JG3 (run2) | 42,301 | 306 | 9,176 | 4,807 | 90,470 | 11,741 | 63 |  |
| *Av* CIP107763<sup>T</sup> | 200,362 | 645 | 1,629 | 1,299 | 98,351 | 7,545 | 135 | 110,391 |
| *Ah* CA-13-1 | 136,486 | 222 | 1,629 | 808 | 62,567 | 2,840 | 46 | 65,195 |
| *Fs* ARS-166-14 | 53,171 | 289 | 5,442 | 1,583 | 1,149,252 | 18,107 | 90 | 36,648 |
| *Fc* FC-100715-19 (run1) | 39,376 | 146 | 3,709 | 836 | 84,881 | 17,593 | 45 | 45,194* |
| *Fc* FC-100715-19 (run2) | 31,121 | 187 | 5,996 | 1,137 | 157,214 | 26,227 | 58 |  |
| *Fc* FC-08-1215-1 | 39,938 | 236 | 5,908 | 1,252 | 106,525 | 22,063 | 74 | 26,486 |

\* indicates the combined total of both runs for that strain

### Quality assessment

The contiguity and quality of each genome assembly was assessed using Quast (v.4.6.3) [57]. Because we lacked reference genomes for comparison, we instead assessed the quality of our genomes using two strategies, focusing on the Pseudonocardia genomes for detailed comparison. First, we compared all genome assemblies to each other based on their shared kmer composition using Mash (v2.0) [58]. These Mash distances were used to construct a phylogeny using Mashtree (v.0.33, available at https://github.com/lskatz/mashtree). Second, we aligned each assembly to their respective Canu+Pilon assembly using MUMmer (v3.1) [59] to identify SNPs and indels relative to the Canu+Pilon assembly. We selected the Canu+Pilon assemblies as references because of their high contiguity and error profiles that were similar to the MiSeq assemblies. However, we stress that this does not comprise a “gold standard” comparison and the relative nature of these comparisons.

### Biosynthetic gene cluster prediction

Secondary metabolite biosynthetic gene clusters (BGCs) were annotated in each Ps JKS002128 assembly using antiSMASH (v.4.1.0) [60]. Fragmented BGCs were annotated by their occurring at contig ends. This likely overestimates the number of fragmented BGCs due to antiSMASH’s tendency to conservatively extend BGCs past their true boundaries. Identical BGCs were identified using the ClustCompare pipeline (available from, https://github.com/klassenlab/ClustCompare). Briefly, PfamScan (v.1.6) [61] was used to annotate protein domains encoded by each BGC and these domains were compared to each other using BLASTp [62]. BGCs were considered to be homologous based on their sharing a minimum ClustCompare similarity score of 0.3 calculated using a 70% similarity threshold between domains in different BGCs, a minimum of two homologous domains shared between BGCs, and a minimum of 50% of the domains in the smaller BGC being homologous to domains in the larger BGC. The resulting homology networks were visualized using Cytoscape (v.3.6.1) [63] to identify clusters of homologous BGCs. Singleton clusters were aligned to the Canu+Pilon genome and individual Canu+Pilon antiSMASH BGCs using MUMmer v3.1 [59] to identify homologies that occurred at the nucleotide level but not at the protein level (e.g., due to high error rates that might confound gene prediction). Nucleotide-level BGC comparisons were also conducted using Mash (v.2.0) [58].

### Insertion Sequence identification

Insertion sequences (ISs) were annotated in the Fs ARS-166-14 Canu, Canu+Pilon, SPAdes, and Unicycler assemblies using ISSaga2 [64]. Full and partial IS sequences were identified by comparing each assembly genome sequence to the ISfinder database. The default detection algorithm and parameters were used for all assemblies in this experiment, and both the total number of hits and those with >70% amino acid similarity to ISs in the ISfinder database were recorded.

## Results

### Sequencing

We sequenced the genomes of nine bacterial strains using both Oxford Nanopore MinION and Illumina MiSeq technologies, together spanning a wide range of GC content (*Flavobacterium*: 31%; *Aeromonas*: 59-61%; *Pseudonocardia*: 74%). MinION sequencing coverage ranged from 40-135X and generated median read lengths of 1,629-9,665 bps (Table 2). Median MinION read lengths for Ah CA-13-1 and Av CIP107763^T^ were considerably shorter than for the other MinION libraries due to difficulties in extracting high molecular weight DNA from these strains. Illumina Nextera libraries were sequenced for all *Aeromonas* and *Flavobacterium* strains with coverage ranging from 30-169X (Table 3). Preliminary Nextera libraries were also constructed for the *Pseudonocardia* strains, but these were highly biased and generated extremely fragmented assemblies (1000s of contigs; data not shown). We therefore instead generated Illumina TruSeq PCR-free libraries for these strains, with coverage ranging from 71-246X (Table 3).

**Table 3.** Summary of Illumina sequencing

|  | **Total Raw**<br>**Reads** | **Total Bases**<br>**(Mbps)** | **Coverage**<br>**(fold)** | **Total Reads**<br>**After Filtering** |
| --- | --- | --- | --- | --- |
| ***Ps* JKS002128** | 6,120,982 | 1,536 | 246 | 5,475,000 |
| ***Ps* JKS002072** | 1,766,572 | 443 | 71 | 1,638,060 |
| ***Ps* JKS002056** | 5,038,846 | 1,265 | 203 | 4,736,206 |
| ***Av* JG3** | 1,488,761 | 372 | 79 | 942,391 |
| ***Av* CIP107763<sup>T</sup>** | 566,606 | 142 | 30 | 536,504 |
| ***Ah* CA-13-1** | 950,886 | 238 | 51 | 873,417 |
| ***Fs* ARS-166-14** | 2,164,975 | 541 | 169 | 890,703 |
| ***Fc* FC-100715-19** | 2,072,592 | 518 | 162 | 1,130,797 |
| ***Fc* FC-08-1215-1** | 1,145,425 | 286 | 89 | 987,428 |

### Genome Assembly

Seven assemblies were generated for each strain, four based on MiSeq data either alone (SPAdes, Unicycler) or with MinION data to deconvolute the MiSeq assembly graph (SPAdes-hybrid, Unicycler-hybrid), and three based on MinION data either alone (Canu), polished using the same MinION data (Canu+Nanopolish), or polished using MiSeq data (Canu+Pilon). Both the SPAdes and Unicycler assemblies had the largest number of contigs out of all assemblies generated for each strain (Figure 1). These assemblies also typically had the lowest N50 values compared to the other assemblies. Ah CA-13-1 and Av CIP107763^T^ were exceptions to this trend, likely due to their lower quality MinION libraries. The addition of MinION reads to deconvolute the SPAdes and Unicycler assembly graphs lowered the number of contigs and increased the N50 for all assemblies (Figure 1). This highlights the ability of long MinION reads to resolve genomic repeats that otherwise stymied assembly of these genomes from short reads. Unicycler consistently outperformed SPAdes during hybrid assembly (the only exception being Av CIP107763^T^) but not when assembling MiSeq reads only.

**Figure 1:**
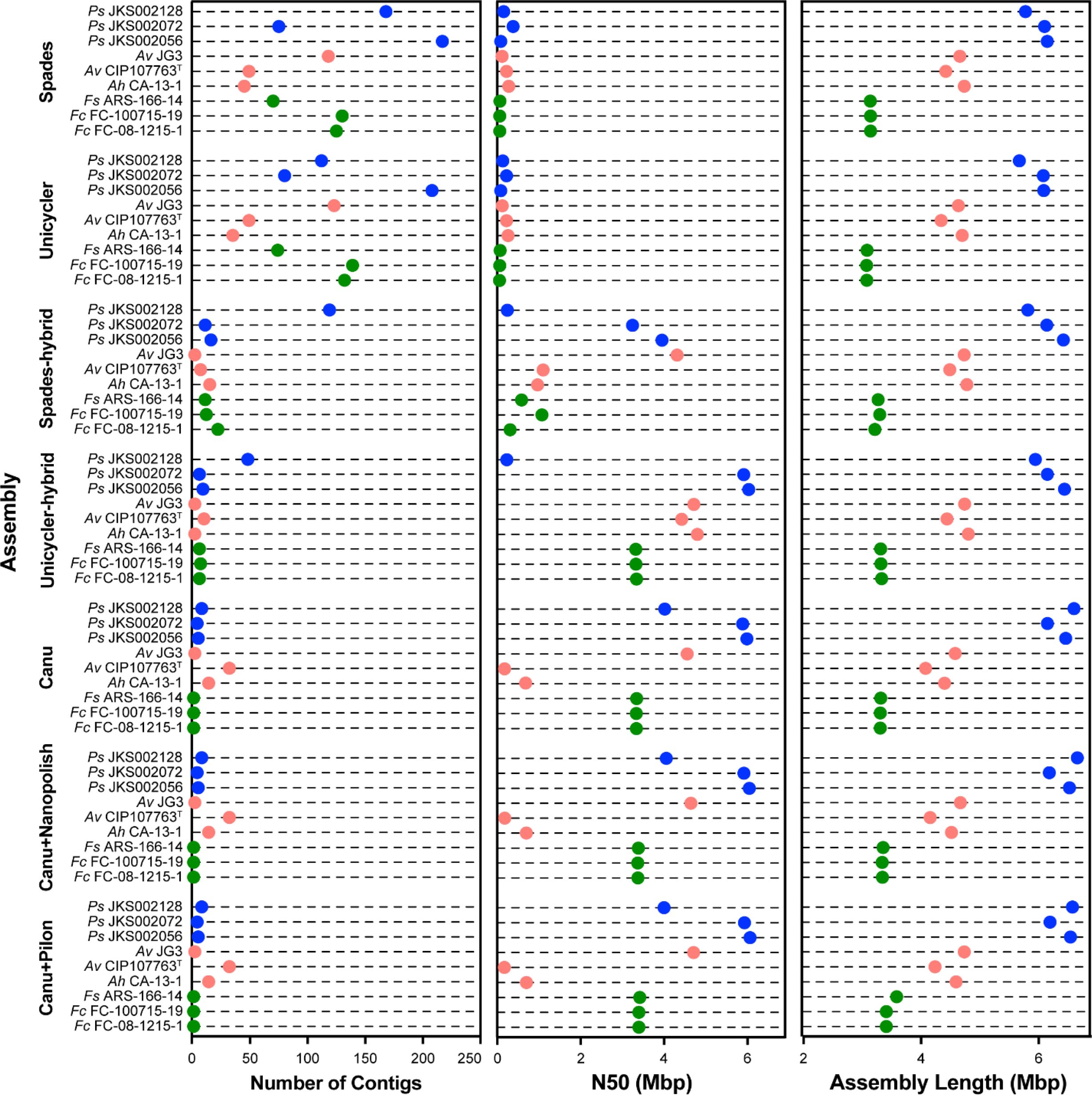
MinION reads improve assembly contiguity. The number of contigs (left), N50 (in Mbp, center), and assembly length (in Mbp, right) are shown for each of the MiSeq-based (SPAdes, Unicycler, SPAdes-hybrid, and Unicycler-hybrid) and MinION-based (Canu, Canu+Nanopolish, Canu+Pilon) genome assemblies. Results for *Pseudonocardia*, *Aeromonas*, and *Flavobacterium* are shown in blue, red, and green, respectively.

Canu assemblies were more contiguous and had higher N50 values than all MiSeq-based assemblies, except for Av CIP107763^T^ Unicycler-hybrid and SPAdes-hybrid assemblies and the Ah CA-13-1 Unicycler-hybrid assembly (Figure 1A, B). These two strains had lower quality MinION libraries (Table 2) that likely compromised the Canu assemblies, even if they were still more contiguous than the MiSeq-only SPAdes and Unicycler assemblies. Canu assemblies were used as the base for polishing with either Nanopolish or Pilon, and so the number of contigs was the same for the Canu, Canu+Nanopolish, and Canu+Pilon assemblies (Figure 1). The Canu assembly sizes were greater than those of any MiSeq-based assembly for all *Flavobacterium* and *Pseudonocardia* strains (up for ~14% for Ps JKS002128; Figure 1), likely reflecting the MinION’s ability to overcome biases in the Illumina libraries for these genomes with low (31%) and high (74%) GC content, respectively. This was not true for the *Aeromonas* assemblies, likely reflecting fewer biases in the Illumina libraries for these strains with more moderate GC content (59-61%). Taken together, these assemblies demonstrate that MinION sequencing improves assembly contiguity, especially where Illumina sequencing libraries are the most biased.

### Assembly Accuracy

Because we lacked high-quality reference genomes for our strains, we instead used several comparative analyses to assess the accuracy of our assemblies. We focused on *Pseudonocardia* for these analyses because these appeared to be the most challenging to assemble based on the substantial differences in their assembly sizes and contiguities (Figure 1). We used Mash [58] to compare all of our *Pseudonocardia* assemblies to each other according to their shared k-mer content and to construct a distance-based phylogeny (Figure 2). Canu assemblies were the least similar to the MiSeq-based assemblies, followed by the Canu+Nanopolish assemblies. This suggests that MinION data alone cannot produce accurate *Pseudonocardia* assemblies using current technologies. These data might alternatively be interpreted to mean that the MiSeq-based assemblies have lower accuracy compared to the Canu and Canu+Nanopolish assemblies, but we consider this unlikely based on previous research that argues against this interpretation [17, 27, 37, 38]. Canu+Pilon assemblies were more similar to the MiSeq-based assemblies, suggesting that polishing MinION-based assemblies with MiSeq reads is an effective strategy to generate microbial genome assemblies that are both accurate and contiguous. However, some divergence was observed between the Canu+Pilon and MiSeq-based genome assemblies. This was especially true for Ps JKS002128, which appeared to have the most biased MiSeq library in our study based on differences in the sizes of the MiSeq-based and MinION-based assemblies for this strain (Figure 1). These differences are consistent with the existence of regions in the Canu assembly that lacked mapping MiSeq reads, leaving these regions uncorrected [65]. All genome assemblies for the same strain clustered together in the Mashtree analysis (Figure 2), indicating that even the high error rates of the Canu and Canu+Nanopolish assemblies did not obscure strain-level phylogenetic differences.

**Figure 2:**
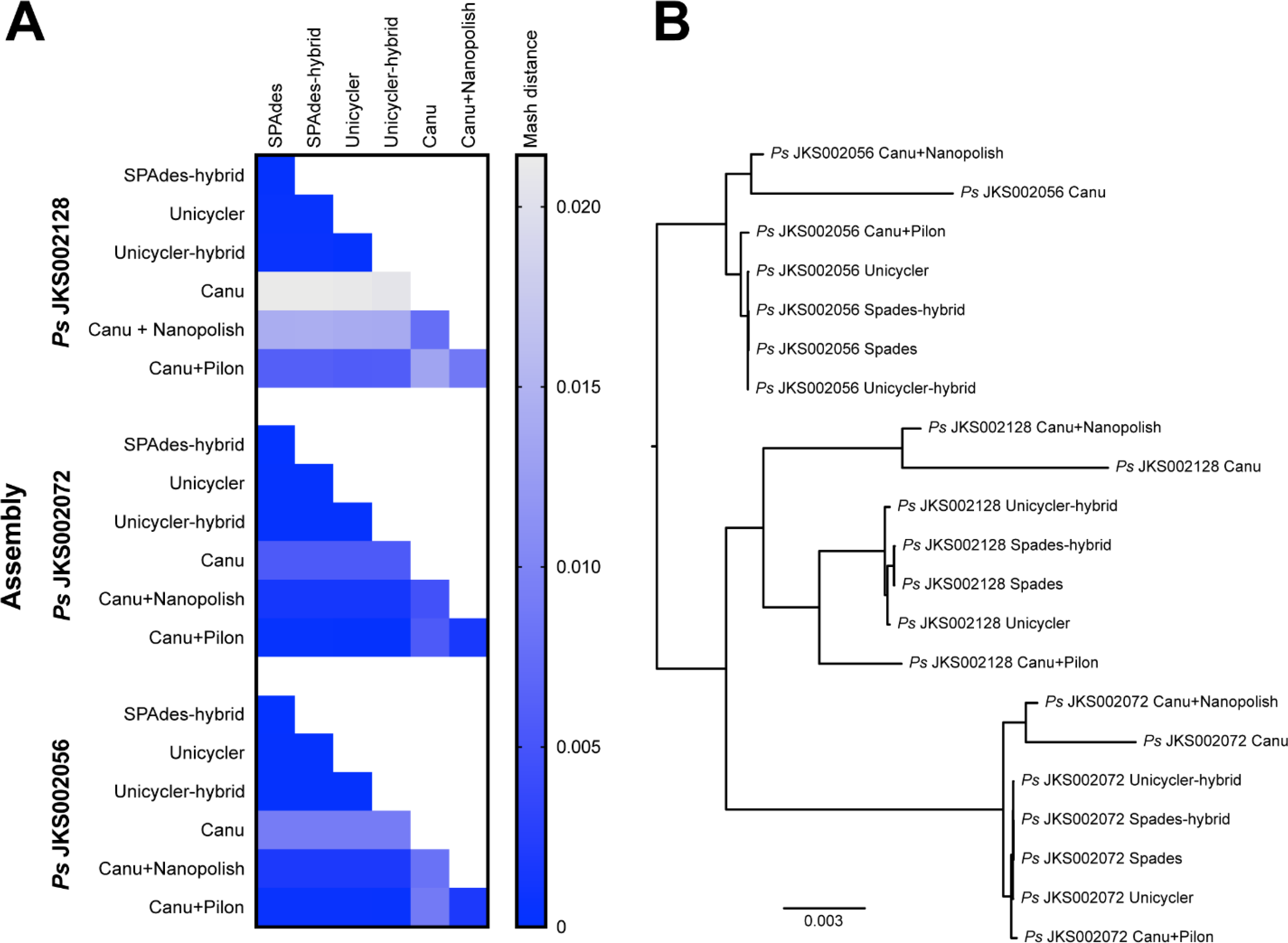
Comparison of Pseudonocardia assemblies generated during this study. (A): Heatmaps depicting Mash distances between the assemblies of each *Pseudonocardia* strain based on their shared k-mer content. Whiter colors indicate greater Mash distances between assemblies. (B): Mashtree analysis showing the relationships of all *Pseudonocardia* assemblies to each other, based on Mash distances. The scale bar represents a Mash distance of 0.003.

Based on the Mash analysis, the Canu+Pilon assemblies were used as a reference against which to compare the other assemblies based on their higher contiguity and substantial accuracy. The high accuracy of MiSeq sequencing meant that all MiSeq-based assemblies had few SNPs and Indels relative to the Canu+Pilon assembly (Figure 3). In contrast, the Canu assemblies had many more SNPs and indels relative to the Canu+Pilon assembly, especially for Ps JKS002056 (Figure 3). Polishing these Canu assemblies using Nanopolish reduced the number of indels, and the number of SNPs to a lesser extent (Figure 3). However, the numbers of SNPs and indels were still much higher than for the MiSeq-based assemblies.

**Figure 3:**
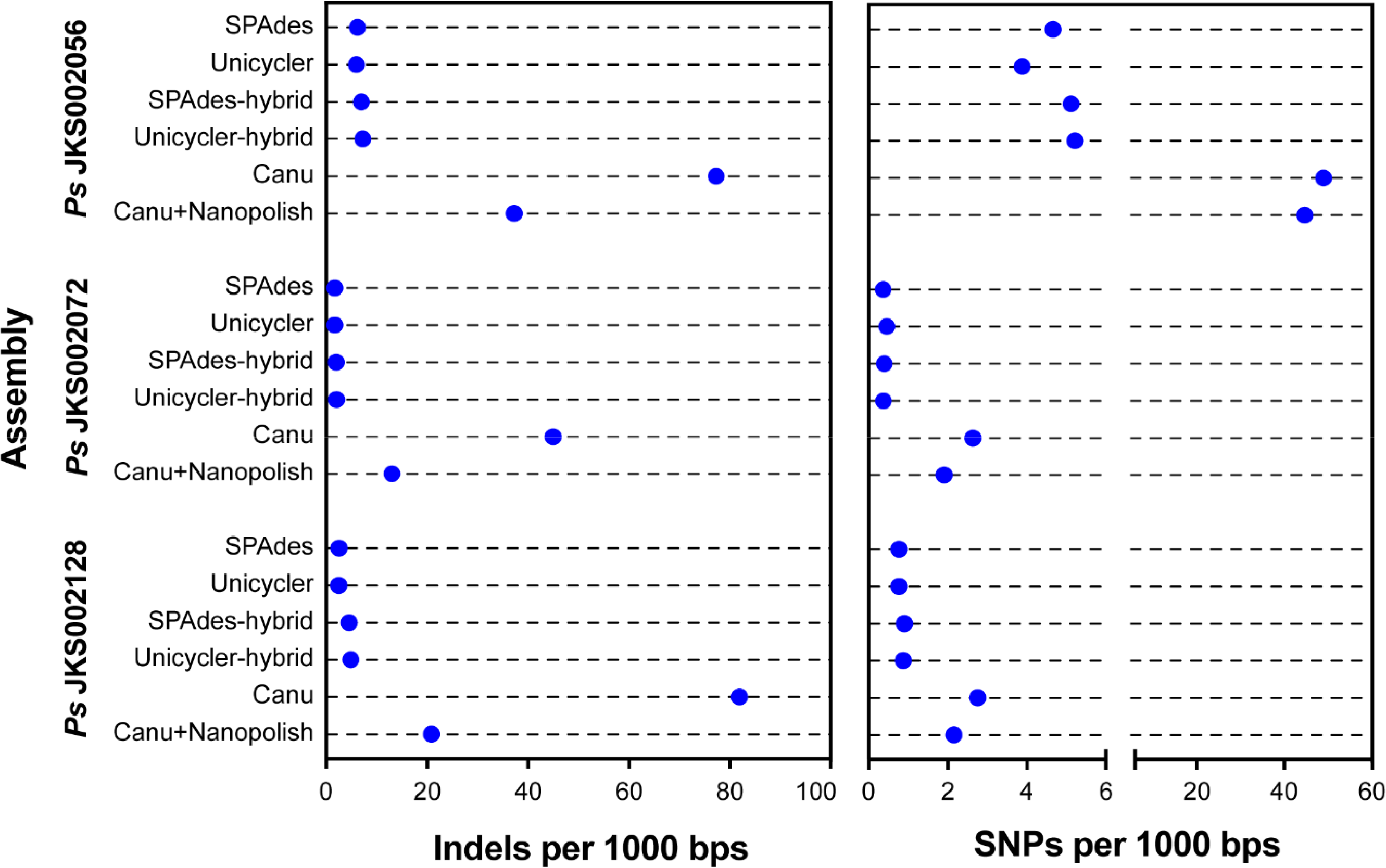
Quantification of insertion/deletions (indels, left) and single nucleotide polymorphisms (SNPs, right) in all *Pseudonocardia* strains sequenced during this study, as determined by aligning each assembly to the Canu+Pilon assembly for that strain as a reference.

### MinION Sequencing Depth

Canu assemblies were performed using 5-7 different levels of coverage for strains Av JG3, Fs ARS-166-14, and Ps JKS002128. These assemblies suggest that the amount of coverage needed for a high-quality MinION-based genome assembly is relatively low, but also depends somewhat on the complexity of each genome. Assemblies for strains Av JG3 and Fs ARS-166-14 did not improve substantially above 30X coverage, consistent with previous findings [66]. However, assemblies for strain Ps JKS002128 improved incrementally up to 70X coverage (Figure 4), suggesting that higher coverage may be necessary for genomes with high GC content. Even though they were assembled into a few contigs, these assemblies were not error-free based on the different genome sizes and N50 values obtained for assemblies using different high-coverage datasets. The single 50X Av JG3 assembly also lacked a plasmid that was present in assemblies for the lower coverage datasets (data not known). Researchers should therefore assess their goals for MinION sequencing before progressing with a run and consider stopping data collection at a certain threshold to conserve flow cells and to decrease sequencing time and cost.

**Figure 4:**
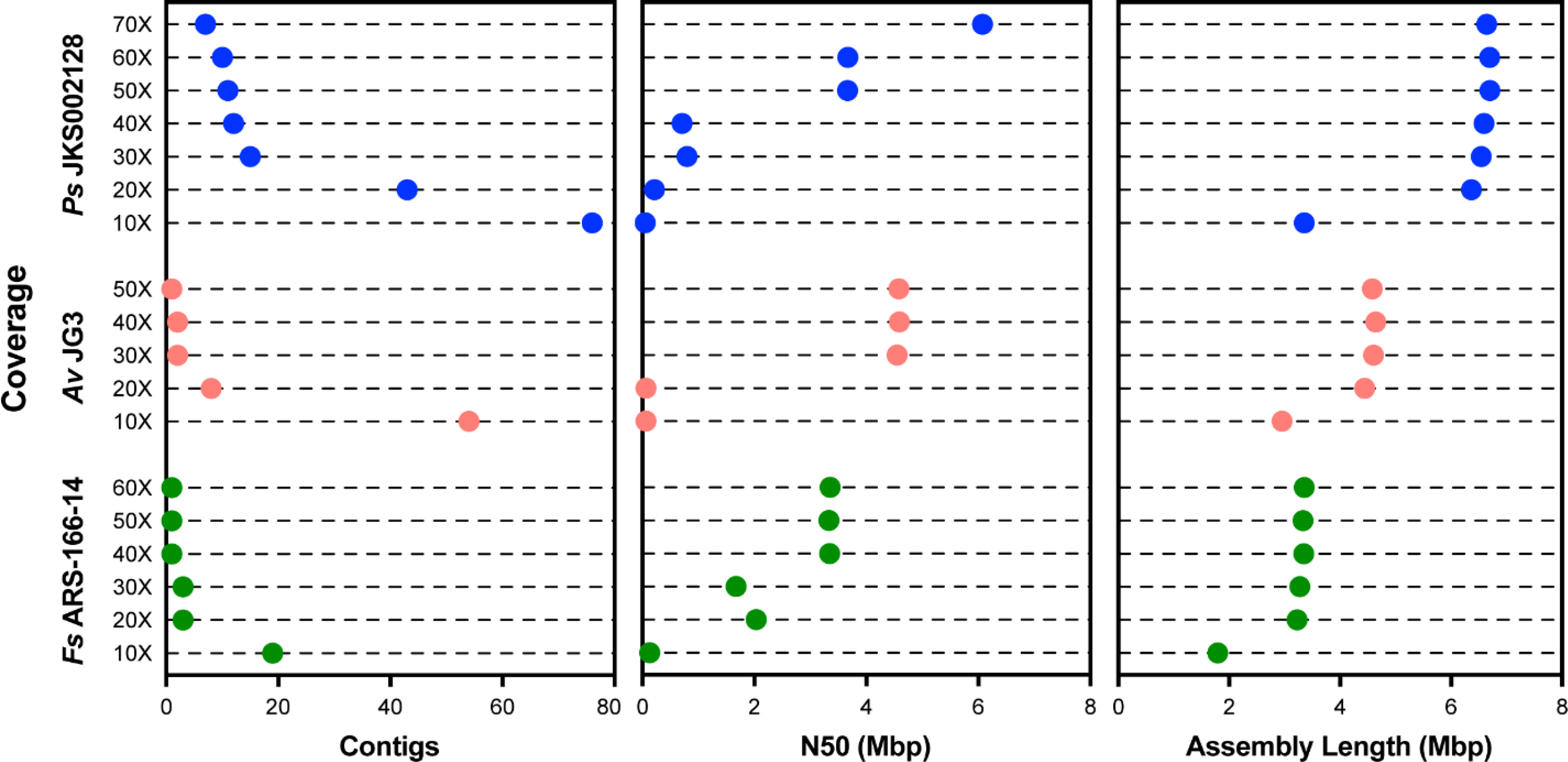
The effect of coverage on Canu genome assembly contiguity. The number of contigs (Left), N50 (in Mbp, Center), and assembly length (in Mbp, Right) are shown for subsets of the Ps JKS002128 (blue), Av JG3 (red), and Fs ARS-166-14 (green) MinION reads used in Figure 1.

### Biosynthetic gene cluster prediction

One expected benefit of high quality genome assemblies is that they will substantially improve the annotation of repetitive genomic regions relative to lower quality assemblies. To test this, we compared antiSMASH [60] secondary metabolite biosynthetic gene cluster (BGC) annotations for all of our Ps JKS002128 assemblies. Actinobacteria such as *Pseudonocardia* typically possess many BGCs, although they are often difficult to assemble correctly [16]. AntiSMASH consistently predicted 12 and 13 BGCs for the SPAdes and Unicycler assembles, respectively, and 12 BGCs for both the SPAdes-hybrid and Unicycler-hybrid assemblies (Figure 5). The extra BGC in the Unicycler assembly is due to there being two separate fragments of BGC 1 annotated in this assembly. More BGCs were predicted for the Canu (17), Canu+Nanopolish (19), and Canu+Pilon (18) assemblies, including 4 BGCs that were found in at least two of these genomes but not in any of the MiSeq-based genomes (Figure 5A). These BGCs may lie at particularly repetitive or bias-prone regions of the Ps JKS002128 genome such that they are omitted from MiSeq-based assemblies but present in MinION-based assemblies that are much less sensitive to these issues. Despite their greater contiguity, the Canu, Canu+Nanopolish, and Canu+Pilon assemblies lacked some combination of BGCs 1, 9, 12, and 13, all of which were found in all of the MiSeq-based assemblies (Figure 5A). The Canu assembly lacked all 4 of these BGCs, the Canu+Nanopolish assembly lacked BGCs 9, 12, and 13, and the Canu+Pilon assembly only lacked BGC 13. These omissions are likely due to gene prediction errors that decreased the ability of antiSMASH to detect these BGCs. Such errors may have also been responsible for the prediction of BGCs 18 and 19 solely in the Canu and Canu+Nanopolish assemblies (Figure 5A), which are likely false positive annotations based on these BGCs only appearing in individual error-prone assemblies. MinION-based genome assemblies therefore substantially increase the sensitivity of BGC annotation, but require polishing to limit annotation errors.

**Figure 5:**
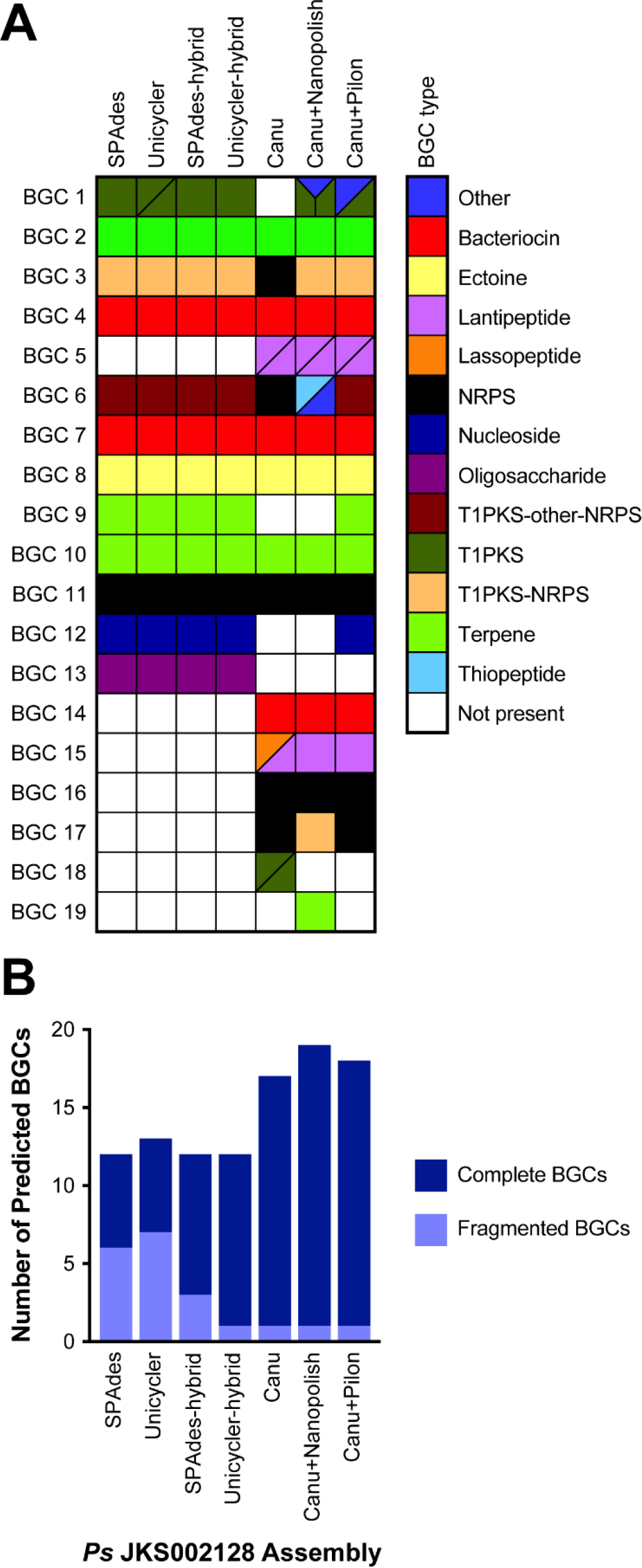
Ps JKS002128 genome assembly quality affects secondary metabolite biosynthetic gene cluster annotation. (A) Homologies between BGCs predicted for each Ps JKS002128 assembly, with each row representing a unique BGC in the Ps JKS002128 genome. Filled boxes indicate the BGCs found in each assembly, colored according to the type of secondary metabolite that it is predicted to encode. White boxes indicate BGCs that were not found in that assembly. Some BGCs occur on multiple contigs or are separated into multiple gene clusters on the same assembly, indicated by either two or three polygons within a single box. BGCs may still be fragmented even if represented by a single box. (B) The total number of complete and fragmented BGCs predicted in each Ps JKS002128 genome assembly.

Improved genome assembly also reduced the number of BGCs that were fragmented, i.e., that overlapped with a contig end (Figure 5B). Approximately half of all BGCs in the SPAdes and Unicycler assemblies were fragmented, reflecting the inability of short-read Illumina data to resolve these repetitive genomic regions. The Unicycler hybrid, and to a lesser extent the SPAdes hybrid, assemblies produced fewer fragmented BGCs, reflecting the increased contiguity of these assemblies. The Canu, Canu+Nanopolish, and Canu+Pilon assemblies all had very few fragmented BGCs, based on BGCs overlapping with contig ends. MinION-based genome assemblies therefore do not only increase the frequency of BGC detection, but also more completely assemble these BGCs and thus increase their value for genome-guided drug discovery. The Canu, Canu+Nanopolish, and Canu+Pilon assemblies did have several annotated gene clusters that were aggregated into a single BGC in other assemblies (Figure 5A). Whether these represent single BGCs that were fragmented in the MinION-based assemblies or multiple BGCs that were located adjacent to each other on the Ps JKS002128 genome is difficult to predict computationally.

### Insertion Sequence Prediction

To further investigate the effect of genome assembly on the annotation of repetitive genetic regions, insertion sequences were predicted in the Fs ARS-166-14 Canu, Canu+Pilon, SPAdes, and Unicycler assemblies using ISSaga2 and the ISfinder database [64]. The total number of full or partial hits to the ISfinder database and the number of hits with amino acid sequence similarities >70% are reported in Figure 6. The Canu+Pilon assembly had the most unique insertion sequences with 70% or greater sequence similarity to the ISfinder database (20), followed by the Canu assembly with 15, and then the Unicycler and SPAdes assemblies with 4 and 3, respectively. Interestingly, the Canu+Pilon assembly also had the greatest total number of hits, but these likely contain many false positive results that require further curation.

**Figure 6:**
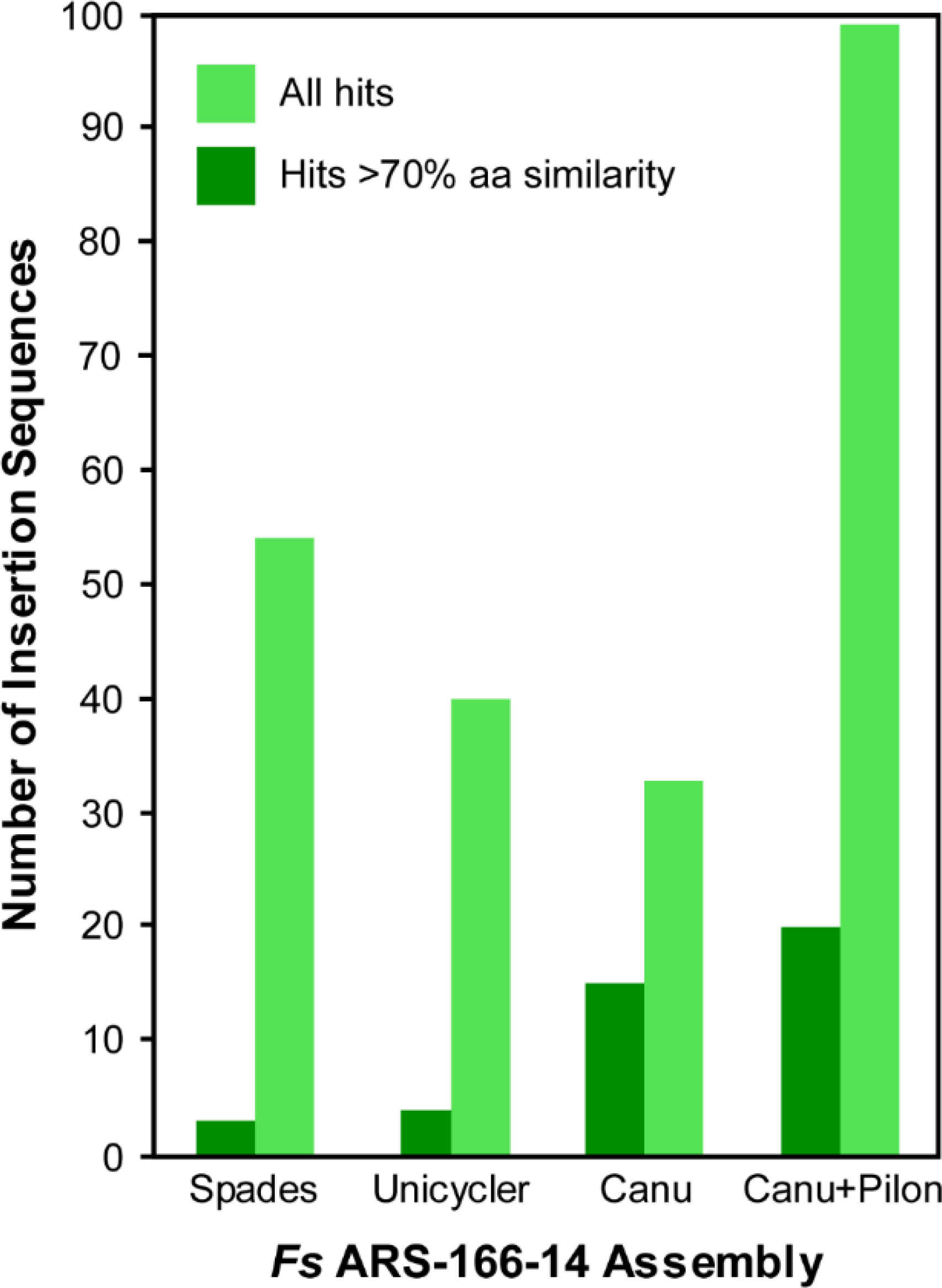
Fs ARS-166-14 genome assembly quality affects insertion sequences annotation. Both the total number of hits and hits with >70% amino acid identity to insertion sequences in the ISfinder database are shown. The former likely includes false-positive annotations while the latter is more conservative.

## Discussion

Single-molecule, long-read sequencing technologies such as the Oxford Nanopore MinION have strong potential to revolutionize the sequencing and de novo assembly of bacterial genomes. Existing short-read sequencing technologies frequently produce genome assemblies that are broken into 10s-100s of contigs, such as in our assemblies generated using only short-read MiSeq data (Figure 1). Fragmented genome assemblies prevent accurate annotation of important genome features such as insertion sequences and secondary metabolite biosynthetic gene clusters (Figures 5 and 6). Technological improvements are therefore necessary to fully understand and exploit these genomic features to cure disease and foster biotechnology.

One key reason for poor genome assembly is the inherently limited length of short-reads. By increasing the read length, long-read sequencing technologies such as the MinION disambiguate genomic repeats and generate fewer contig breaks (e.g., [38]). This was clearly evident from our SPAdes- and Unicycler-hybrid assemblies, where the long MinION reads were able to deconvolute the assembly graph produced from the MiSeq data and yielded fewer and longer contigs compared to the MiSeq-only assemblies (Figure 1). Such improvements are likely to continue as MinION-compatible extraction methods for high-molecular weight DNA are refined.

However, this approach assumes that the entire genome is represented in the Illumina sequencing graph, which may not be true because of biases in short-read sequencing library preparation. As a result, some regions of the genome are sequenced to low coverage or excluded entirely, resulting in assembly fragmentation due to missing data. These problems include PCR biases against extreme %GC sequences [8–12] and due to biased insertion of transposases during library preparation [14]. Reflecting such biases, our initial *Pseudonocardia* sequencing experiments that used the Illumina Nextera library preparation method (which includes both transposases and PCR) produced genome assemblies with 1,000s of contigs (data not shown), compared to the 10s-100s of *Pseudonocardia* contigs produced using Illumina TruSeq PCR-free libraries (Figure 1). Single-molecule sequencing methods such as the MinION avoid many of these biases by sequencing individual template DNA molecules without using PCR. This is reflected by the higher contiguity of our *Pseudonocardia* Canu genome assemblies compared to the SPAdes- and Unicycler-hybrid assemblies that used MinION reads to deconvolute the potentially biased Illumina assembly graphs (Figure 1). All of our *Flavobacterium* and *Pseudonocardia* Canu assemblies are also larger than those based on Illumina reads, reflecting the inclusion of sequences that were missing from the Illumina sequencing libraries. For *Pseudonocardia*, these differences were sometimes substantial (up to a 13.7% increase in genome size). These results point to library preparation bias as a second source of error common to short-read sequencing that can be overcome by long-read, single-molecule sequencing technologies such as the MinION, in addition to the ability of MinION reads to span long genomic repeats.

Our results also highlight the importance of efficient high molecular weight DNA extraction methods for MinION sequencing. Of the 9 genomes that we sequenced during this study, the two with the lowest median read length (Ah CA-13-1 and Av CIP107763^T^) produced the least contiguous Canu assemblies (14 and 32 contigs, respectively). However, this is still more contiguous than the MiSeq-only SPAdes and Unicycler assemblies for these strains. MinION reads also improved these SPAdes and Unicycler assemblies when run in hybrid mode, demonstrating the utility of long reads even if DNA extraction remains suboptimal. There is a current need for reliable protocols to produce high molecular weight genomic DNA that is compatible with the MinION sequencer, and the Oxford Nanopore Voltrax and Ubik devices (https://nanoporetech.com/about-us/news/clive-g-brown-cto-plenary-london-calling) show strong potential to overcome these issues. The degree to which such devices are compatible with diverse cell wall chemistries remains to be validated.

Although most of our MinION-based assemblies were more contiguous than the MiSeq-based assemblies, they were less accurate. Assemblies generated using Canu contained a large number of SNP and indels relative to our Illumina-based assemblies (Figures 2 and 3). These differences were reduced by using Nanopolish to correct the Canu assembly using MinION reads, and even better results were obtained using Pilon to correct the Canu assembly using MiSeq reads (Figures 2 and 3). However, differences still existed between these polished assemblies and the Illumina assemblies in some cases (most obviously for Pseudonocardia sp. JKS002128). Although it is possible that the MiSeq assemblies contained errors relative to the MinION assemblies, this would be inconsistent with previous work comparing MinION assemblies to high-quality reference genomes [17, 27, 37, 38]. Illumina reads are also unable to correct repetitive genome sequences that cannot be unambiguously mapped using short reads, and so these regions will be uncorrected even in Canu+Pilon assemblies [65]. A tradeoff therefore exists between the higher contiguity of MinION-based assemblies relative to their higher number of SNP and indel errors. Minimizing such errors is a current technological focus of ONT (https://nanoporetech.com/about-us/news/clive-g-brown-cto-plenary-london-calling) and so this tradeoff may lessen in the near future.

The importance of these assembly trade-offs is highlighted by our analysis of repetitive genomic regions. For example, antiSMASH annotated ~1/3 more secondary metabolite biosynthetic gene clusters (BGC) in the MinION-based assemblies of *Pseudonocardia* sp. JKS002128 compared to the MiSeq-based assemblies (Figure 5), confirming our previous observations that BGCs are poorly resolved by Illumina sequencing [16]. Similar results were obtained when annotating insertion sequences in *Flavobacterium* sp. Fs ARS-166-14, as expected due to the highly repetitive nature of these genomic regions (Figure 6). The BGCs that were annotated in the Illumina-only assemblies were highly fragmented, highlighting the challenge of sequencing these complex genomic regions (Figure 5). Interestingly, the genome assemblies that contained the highest number of SNP and indel errors (Figure 2) contained several BGCs that were unique to those particular genomes (Figure 5), and lacked several BGCs that were annotated in the MiSeq-based assemblies. These differences are likely due to the difficulty in accurately predicting gene structures in highly error-prone genomes due to gene truncation or misplaced start sites. Indeed, our initial ClustCompare analysis to compare BGCs based on their protein sequences did not detect many true homologies between BGCs annotated in the Canu and Canu+Nanopolish assemblies to those annotated in assemblies that were generated or polished using MiSeq data due to the large number of misannotated gene structures in the Canu and Canu+Nanopolish assemblies (data not shown). These homologies only became clear using comparisons between nucleotide sequences. High numbers of SNP and indel errors can therefore prevent accurate genome annotation due to errors in gene structure prediction. Several homologous BGCs were also annotated as having different biosynthetic classes in different genomes (represented by the different colors in Figure 5). Together, these analyses highlight the importance of contiguous and accurate genome assemblies for the prediction of repetitive elements such as BGCs, and highlight the utility of MinION sequencing in this application, especially when polished using accurate Illumina reads.

In summary, our data highlights the ability of long-read, single-molecule MinION sequencing to overcome current limitations of short-read sequencing, particularly its inability to disambiguate repetitive genome regions and avoid biases introduced during library preparation. Overcoming these limitations greatly improves the annotation of many clinically-and biotechnologically-important genomic regions such as insertion sequences and BGCs (Figures 5 and 6). However, SNP and indel errors remain problematic in de novo assemblies generated from MinION data. This is likely to improve in the near future given the extensive research underway in this area. Because twelve microbial genomes can currently be sequenced to sufficient coverage (40-50X; Figure 3) on a single MinION or MiSeq flowcell, combining these data currently requires ~$100-$200 for the MinION and ~$150 for Illumina sequencing in reagent and consumable costs per genome. Combining these two data types is therefore an affordable means to dramatically increase the quality of any bacterial de *novo* genome assembly, regardless of their genome complexity or %GC content, and compares favorably to the cost of PacBio sequencing. Future technical advances will likely decrease these costs further, and we anticipate that highly contiguous and accurate de *novo* assembly of bacterial genomes will become standard in the field in the very near future.

## Acknowledgements

Funding for this work was provided by NSF IOS-1656475 and the University of Connecticut (J. L.K.) and by USDA-ARS agreement 58-1930-4-002 (J. G.). We thank the UConn Microbiology Analysis, Resources, and Services (MARS) and Center for Genomic Information (CGI) facilities for assistance with the Illumina sequencing and Susan Janton for excellent technical assistane in preparing some of the Illumina libraries. We also thank the Florida, Georgia, and North Carolina Departments of Environmental Protection for permission and assistance with our ant collections on state lands, and Drs. Greg Wiens and Tim Welch from the USDA National Center for Cool and Cold Water Aquaculture, Agriculture Research Service, Kearneysville, West Virginia, USA for providing the *Flavobacterium* strains.

## References

1. Goodwin S, McPherson JD, McCombie WR. Coming of age: ten years of next-generation sequencing technologies. Nat Rev Genet. 2016;17:333–51. doi:10.1038/nrg.2016.49.

2. Shendure J, Balasubramanian S, Church GM, Gilbert W, Rogers J, Schloss JA, et al. DNA sequencing at 40: past, present and future. Nature. 2017;550:345–53. doi:10.1038/nature24286.

3. Whiteford N, Haslam N, Weber G, Prügel-Bennett A, Essex JW, Roach PL, et al. An analysis of the feasibility of short read sequencing. Nucleic Acids Res. 2005;33:e171. doi:10.1093/nar/gni170.

4. Haubold B, Wiehe T. How repetitive are genomes? BMC Bioinformatics. 2006;7:541.

5. Kingsford C, Schatz MC, Pop M. Assembly complexity of prokaryotic genomes using short reads. BMC Bioinformatics. 2010;11:21. doi:1471-2105-11-21 [pii] 10.1186/1471-2105-11-21.

6. Cahill MJ, Köser CU, Ross NE, Archer JAC. Read length and repeat resolution: Exploring prokaryote genomes using next-generation sequencing technologies. PLoS One. 2010;5:e11518.

7. Koren S, Harhay GP, Smith TP, Bono JL, Harhay DM, Mcvey SD, et al. Reducing assembly complexity of microbial genomes with single-molecule sequencing. Genome Biol. 2013;14:R101. doi:10.1186/gb-2013-14-9-r101.

8. Chen YC, Liu T, Yu CH, Chiang TY, Hwang CC. Effects of GC bias in next-generation-sequencing data on de novo genome assembly. PLoS One. 2013;8:e62856.

9. Cheung M-S, Down TA, Latorre I, Ahringer J. Systematic bias in high-throughput sequencing data and its correction by BEADS. Nucleic Acids Res. 2011;39:e103. doi:10.1093/nar/gkr425.

10. Benjamini Y, Speed TP. Summarizing and correcting the GC content bias in high-throughput sequencing. Nucleic Acids Res. 2012;40:e72. doi:10.1093/nar/gks001.

11. Aird D, Ross MG, Chen W-S, Danielsson M, Fennell T, Russ C, et al. Analyzing and minimizing PCR amplification bias in Illumina sequencing libraries. Genome Biol. 2011;12:R18. doi:10.1186/gb-2011-12-2-r18.

12. Marine R, Polson SW, Ravel J, Hatfull G, Russell D, Sullivan M, et al. Evaluation of a transposase protocol for rapid generation of shotgun high-throughput sequencing libraries from nanogram quantities of DNA. Appl Environ Microbiol. 2011;77:8071–9.

13. Muto A, Osawa S. The guanine and cytosine content of genomic DNA and bacterial evolution. Proc Natl Acad Sci U S A. 1987;84:166–9.

14. Lan JH, Yin Y, Reed EF, Moua K, Thomas K, Zhang Q. Impact of three Illumina library construction methods on GC bias and HLA genotype calling. Hum Immunol. 2015;76:166–75. doi:10.1016/j.humimm.2014.12.016.

15. Acuña-Amador L, Primot A, Cadieu E, Roulet A, Barloy-Hubler F. Genomic repeats, misassembly and reannotation: a case study with long-read resequencing of *Porphyromonas gingivalis* reference strains. BMC Genomics. 2018;19:54.

16. Klassen JL, Currie CR. Gene fragmentation in bacterial draft genomes: extent, consequences and mitigation. BMC Genomics. 2012;13:14.

17. Sović I, Križanović K, Skala K, Šikić M. Evaluation of hybrid and non-hybrid methods for de novo assembly of nanopore reads. Bioinformatics. 2016;32:2582–9.

18. Fraser CM, Eisen JA, Nelson KE, Ian T, Salzberg SL, Paulsen IT. The value of complete microbial genome sequencing (you get what you pay for). J Bacteriol. 2002;184:6403–5. doi:10.1128/JB.184.23.6403.

19. Mardis E, McPherson J, Martienssen R, Wilson RK, McCombie WR. What is finished, and why does it matter. Genome Res. 2002;12:669–71.

20. Leggett RM, Clark MD. A world of opportunities with nanopore sequencing. J Exp Bot. 2017;68:5419–29.

21. Quick J, Loman NJ, Duraffour S, Simpson JT, Severi E, Cowley L, et al. Real-time, portable genome sequencing for Ebola surveillance. Nature. 2016;530:228–32. doi:10.1038/nature16996.

22. Payne A, Holmes N, Rakyan V, Loose M. Whale watching with BulkVis: A graphical viewer for Oxford Nanopore bulk fast5 files. bioarXiv. 2018;:312256.

23. Ip CLC, Loose M, Tyson JR, de Cesare M, Brown BL, Jain M, et al. MinION Analysis and Reference Consortium: Phase 1 data release and analysis. F1000Research. 2015;4:1075. doi:10.12688/f1000research.7201.1.

24. Jain M, Tyson JR, Loose M, Ip CLC, Eccles DA, O’Grady J, et al. MinION Analysis and Reference Consortium: Phase 2 data release and analysis of R9.0 chemistry. F1000Research. 2017;6:760. doi:10.12688/f1000research.11354.1.

25. Lu H, Giordano F, Ning Z. Oxford Nanopore MinION sequencing and genome assembly. Genomics Proteomics Bioinformatics. 2016;14:265–79. doi:10.1016/j.gpb.2016.05.004.

26. Senol Cali D, Kim JS, Ghose S, Alkan C, Mutlu O. Nanopore sequencing technology and tools for genome assembly: computational analysis of the current state, bottlenecks and future directions. Brief Bioinform. 2018. doi:10.1093/bib/bby017.

27. Magi A, Semeraro R, Mingrino A, Giusti B, D’Aurizio R. Nanopore sequencing data analysis: state of the art, applications and challenges. Brief Bioinform. 2017. doi:10.1093/bib/bbx062.

28. de Lannoy C, de Ridder D, Risse J. A sequencer coming of age: *de novo* genome assembly using MinION reads. F1000Research. 2017;6:1083. doi:10.12688/f1000research.12012.1.

29. Ashton PM, Nair S, Dallman T, Rubino S, Rabsch W, Mwaigwisya S, et al. MinION nanopore sequencing identifies the position and structure of a bacterial antibiotic resistance island. Nat Biotechnol. 2015;33:296–300.

30. Risse J, Thomson M, Patrick S, Blakely G, Koutsovoulos G, Blaxter M, et al. A single chromosome assembly of *Bacteroides fragilis* strain BE1 from Illumina and MinION nanopore sequencing data. Gigascience. 2015;4:60. doi:10.1186/s13742-015-0101-6.

31. Karlsson E, Lärkeryd A, Sjödin A, Forsman M, Stenberg P. Scaffolding of a bacterial genome using MinION nanopore sequencing. Sci Rep. 2015;5:11996. doi:10.1038/srep11996.

32. Wick RR, Judd LM, Gorrie CL, Holt KE. Unicycler: Resolving bacterial genome assemblies from short and long sequencing reads. PLOS Comput Biol. 2017;13:e1005595.

33. Antipov D, Korobeynikov A, McLean JS, Pevzner PA. hybridSPAdes: an algorithm for hybrid assembly of short and long reads. Bioinformatics. 2016;32:1009–15.

34. Loman NJ, Quick J, Simpson JT. A complete bacterial genome assembled *de novo* using only nanopore sequencing data. Nat Methods. 2015;12:733–5. doi:10.1101/015552.

35. Koren S, Walenz BP, Berlin K, Miller JR, Bergman NH, Phillippy AM. Canu: scalable and accurate long-read assembly via adaptive k-mer weighting and repeat separation. Genome Res. 2017;27:722–36.

36. Walker BJ, Abeel T, Shea T, Priest M, Abouelliel A, Sakthikumar S, et al. Pilon: An integrated tool for comprehensive microbial variant detection and genome assembly improvement. PLoS One. 2014;9:e112963.

37. Judge K, Hunt M, Reuter S, Tracey A, Quail MA, Parkhill J, et al. Comparison of bacterial genome assembly software for MinION data and their applicability to medical microbiology. Microb Genomics. 2016;2. doi:10.1099/mgen.0.000085.

38. George S, Pankhurst L, Hubbard A, Votintseva A, Stoesser N, Sheppard AE, et al. Resolving plasmid structures in Enterobacteriaceae using the MinION nanopore sequencer: assessment of MinION and MinION/Illumina hybrid data assembly approaches. Microb Genomics. 2017;3:doi:10.1099/mgen.0.000118.

39. Wick RR, Judd LM, Gorrie CL, Holt KE. Completing bacterial genome assemblies with multiplex MinION sequencing. Microb Genomics. 2017;3. doi:10.1099/mgen.0.000132.

40. Bayliss SC, Hunt VL, Yokoyama M, Thorpe HA, Feil EJ. The use of Oxford Nanopore native barcoding for complete genome assembly. Gigascience. 2017;6:1–6. doi:10.1093/gigascience/gix001.

41. Todd MS, Settlage RE, Lahmers KK, Slade DJ. *Fusobacterium* genomics using MinION and Illumina sequencing enables genome completion and correction. bioRxiv. 2018;:305573.

42. Laver T, Harrison J, O’Neill PA, Moore K, Farbos A, Paszkiewicz K, et al. Assessing the performance of the Oxford Nanopore Technologies MinION. Biomol Detect Quantif. 2015;3:1–8. doi:10.1016/j.bdq.2015.02.001.

43. Bainomugisa A, Duarte T, Lavu E, Pandey S, Coulter C, Marais B, et al. A complete nanonpore-only assembly of an XDR *Mycobacterium tuberculosis* Beijing lineage strain identifies novel genetic variation in repetitive PE/PPE gene regions. bioRxiv. 2018;:256719. doi:10.1101/256719.

44. Marden JN, McClure EA, Beka L, Graf J. Host matters: medicinal leech digestive-tract symbionts and their pathogenic potential. Front Microbiol. 2016;7:1569.

45. Oh D-C, Poulsen M, Currie CR, Clardy J. Dentigerumycin: a bacterial mediator of an ant-fungus symbiosis. Nat Chem Biol. 2009;5:391–3. doi:10.1038/nchembio.159.

46. Beka L, Fullmer MS, Colston SM, Nelson MC, Talagrand-Reboul E, Walker P, et al. Low-level antimicrobials in the medicinal leech select for resistant pathogens that spread to patients. mBio. 2018;9:e01328–18.

47. Colston SM, Fullmer MS, Beka L, Lamy B, Peter Gogarten J, Graf J. Bioinformatic genome comparisons for taxonomic and phylogenetic assignments using aeromonas as a test case. mBio. 2014;5:e02136–14.

48. Indergand S, Graf J. Ingested blood contributes to the specificity of the *symbiosis of Aeromonas veronii* Biovar Sobria and *Hirudo medicinalis*, the medicinal leech. Appl Environ Microbiol. 2000;66:4735–41.

49. Cain KD, LaFrentz BR. Laboratory Maintenance of *Flavobacterium psychrophilum* and *Flavobacterium columnare*. Curr Protoc Microbiol. 2017;6:13B.1.1–13B.1.12.

50. Marsh SE, Poulsen M, Gorosito NB, Pinto-Tomás A, Masiulionis VE, Currie CR. Association between *Pseudonocardia* symbionts and *Atta* leaf-cutting ants suggested by improved isolation methods. Int Microbiol. 2013;16:17–25.

51. Rio RVM, Anderegg M, Graf J. Characterization of a catalase gene from *Aeromonas veronii*, the digestive-tract symbiont of the medicinal leech. Microbiology. 2007;153:1897–906.

52. Nelson K, Selander RK. Analysis of genetic variation by polymerase chain reaction-based nucleotide sequencing. Methods Enzymol. 1994;235:174–83.

53. Bolger AM, Lohse M, Usadel B. Trimmomatic: a flexible trimmer for Illumina sequence data.Bioinformatics. 2014;30:2114–20.

54. Loman NJ, Quinlan AR. Poretools: a toolkit for analyzing nanopore sequence data. Bioinformatics. 2014;30:3399–401.

55. Li H, Durbin R. Fast and accurate short read alignment with Burrows-Wheeler transform. Bioinformatics. 2009;25:1754–60.

56. Li H, Handsaker B, Wysoker A, Fennell T, Ruan J, Homer N, et al. The sequence alignment/map format and SAMtools. Bioinformatics. 2009;25:2078–9.

57. Gurevich A, Saveliev V, Vyahhi N, Tesler G. QUAST: Quality assessment tool for genome assemblies. Bioinformatics. 2013;29:1072–5.

58. Ondov BD, Treangen TJ, Melsted P, Mallonee AB, Bergman NH, Koren S, et al. Mash: fast genome and metagenome distance estimation using MinHash. Genome Biol. 2016;17:132. doi:10.1186/s13059-016-0997-x.

59. Kurtz S, Phillippy A, Delcher AL, Smoot M, Shumway M, Antonescu C, et al. Versatile and open software for comparing large genomes. Genome Biol. 2004;5:R12.

60. Blin K, Wolf T, Chevrette MG, Lu XH, Schwalen CJ, Kautsar SA, et al. antiSMASH 4.0— improvements in chemistry prediction and gene cluster boundary identification. Nucleic Acids Res. 2017;45:W36–41.

61. Finn RD, Coggill P, Eberhardt RY, Eddy SR, Mistry J, Mitchell AL, et al. The Pfam protein families database: towards a more sustainable future. Nucleic Acids Res. 2017;44:D279–85.

62. Camacho C, Coulouris G, Avagyan V, Ma N, Papadopoulos J, Bealer K, et al. BLAST+: architecture and applications. BMC Bioinformatics. 2009;10:421. doi:10.1186/1471-2105-10-421.

63. Shannon P, Markiel A, Ozier O, Baliga NS, Wang JT, Ramage D, et al. Cytoscape: a software environment for integrated models of biomolecular interaction networks. Genome Res. 2003;13:2498–504.

64. Varani AM, Siguier P, Gourbeyre E, Charneau V, Chandler M. ISsaga is an ensemble of web-based methods for high throughput identification and semi-automatic annotation of insertion sequences in prokaryotic genomes. Genome Biol. 2011;12:R30. doi:10.1186/gb-2011-12-3-r30.

65. Watson M. Mind the gaps - ignoring errors in long read assemblies critically affects protein prediction. bioRxiv. 2018;:285049. doi:10.1101/285049.

66. Giordano F, Aigrain L, Quail MA, Coupland P, Bonfield JK, Davies RM, et al. *De novo* yeast genome assemblies from MinION, PacBio and MiSeq platforms. Sci Rep. 2017;7:3935.

